# A chromosome-level reference genome for the common octopus, *Octopus vulgaris* (Cuvier, 1797)

**DOI:** 10.1101/2023.05.16.540928

**Authors:** Dalila Destanović, Darrin T. Schultz, Ruth Styfhals, Fernando Cruz, Jèssica Gómez-Garrido, Marta Gut, Ivo Gut, Graziano Fiorito, Oleg Simakov, Tyler S. Alioto, Giovanna Ponte, Eve Seuntjens

**Affiliations:** Department of Neurosciences and Developmental Biology, University of Vienna, Djerassiplatz 1, 1030, Vienna, Austria; Lab of Developmental Neurobiology, Animal Physiology and Neurobiology Division, Department of Biology, KU Leuven, Leuven, Belgium; KU Leuven Institute for Single Cell Omics (LISCO), KU Leuven, Leuven, Belgium; Leuven Brain Institute, KU Leuven, Leuven, Belgium; Department of Biology and Evolution of Marine Organisms, Stazione Zoologica Anton Dohrn, Villa Comunale, 80121, Naples, Italy; Centro Nacional de Análisis Genómico (CNAG), C/Baldiri Reixac 4, 08028 Barcelona, Spain

**Author notes:** equal contribution (Dalila Destanović, Darrin T. Schultz).

**Keywords:** coleoid cephalopods, chromosome-scale, Hi-C

## Abstract

Cephalopods are emerging animal models and include iconic species for studying the link between genomic innovations and physiological and behavioral complexities. Coleoid cephalopods possess the largest nervous system among invertebrates, both for cell counts and brain-to-body ratio. *Octopus vulgaris* has been at the center of a long-standing tradition of research into diverse aspects of cephalopod biology, including behavioral and neural plasticity, learning and memory recall, regeneration, and sophisticated cognition. However, no chromosome-scale genome assembly is available for *O. vulgaris* to aid in functional studies. To fill this gap, we sequenced and assembled a chromosome-scale genome of the common octopus, *O. vulgaris*. The final assembly spans 2.8 billion basepairs, 99.34% of which are in 30 chromosome-scale scaffolds. Hi-C heatmaps support a karyotype of 1n=30 chromosomes. Comparisons with other octopus species’ genomes show a conserved octopus karyotype, and a pattern of local genome rearrangements between species. This new chromosome-scale genome of *O. vulgaris* will further facilitate research in all aspects of cephalopod biology, including various forms of plasticity and the neural machinery underlying sophisticated cognition, as well as an understanding of cephalopod evolution.

## Introduction

Coleoid cephalopods (cuttlefish, squid, and octopus) comprise about 800 extant species characterized by highly diversified lifestyles, body plans, and adaptations. Cephalopod-specific traits, such as complex nervous systems (Young 1964; Hochner et al. 2006; Hochner 2012; Fiorito et al. 2014; Wang and Ragsdale 2019; Ponte et al. 2021), advanced learning abilities (reviewed in Marini et al. 2017), and the richness in body patterning considered to be involved in camouflaging and communication (Borrelli et al. 2006; Chiao and Hanlon 2019) have made this taxon ideal for studying evolutionary novelties. The neural plasticity of cephalopod brains and the existence of evidence for functionally analogous structures shared with mammalian brains have made cephalopods into a model comparative clade for neurophysiology research (Shigeno et al. 2018; Styfhals et al. 2022).

Despite the technical difficulties of sequencing their typically large and repetitive genomes, the available cephalopod genomes have given insights into the genomic basis for the evolution of novelty (Albertin et al. 2015, 2022; Jiang et al. 2022; Kim et al. 2018; Li et al. 2020; Marino et al. 2022; Schmidbaur et al. 2022). The first-published cephalopod genome, that of *Octopus bimaculoides* (Albertin et al. 2015), made it clear that cephalopod genomic novelties were not attributable to whole-genome duplication, as occurred in the vertebrate ancestor (Meyer and Schartl 1999; Dehal and Boore 2005). Comparisons of recently available chromosome-scale genome assemblies, including those of the Boston market squid *Doryteuthis pealeii* (Albertin et al. 2022) and the Hawaiian bobtail squid *Euprymna scolopes* (Schmidbauer et al. 2022), have shown the impact of genome reorganization on novel regulatory units in coleoid cephalopods. Still, it is not yet known how these units are made in terms of their gene content or their evolution in separate squid and octopus lineages. In this respect, it is crucial that the growing cephalopod genomics resources and approaches help obtain high-quality genomes for the established experimental species.

The common octopus, *Octopus vulgaris*, has long been used as a model for the study of learning and cognitive capabilities in invertebrates (reviewed in: Young 1964; Marini et al. 2017), and is also used as a comparative system in the study of neural organization and evolution (Shigeno et al. 2018; Ponte et al. 2022). Furthermore, recent advances in the culture of this species’ early life stages have increased its suitability for molecular approaches and have provided important developmental staging information (Deryckere et al. 2020).

One bottleneck to studying *O. vulgaris* is the lack of a chromosome-scale genome assembly. While the reported karyotype of *O. vulgaris* is 1n=28 (Inaba 1959; Vitturi et al. 1982) or 1n=30 (Gao & Natsukari, 1990), to date there is no definitive answer. Existing genomic resources for *O. vulgaris* include a short read-based genome assembly (Zarrella et al. 2019), and a genome annotation based on the closely related *O. sinensis* genome that is supported with PacBio Iso-Seq reads and FLAM-seq curation (Styfhals et al. 2022; Zolotarov et al. 2022). These resources have been valuable in characterizing the molecular and cellular diversity of the developing brain (Styfhals et al. 2022), the evolution of cephalopod brains (Zolotarov et al. 2022), and the non-coding RNA repertoire unique to cephalopods (Petrosino et al. 2022). Further improvements to the *O. vulgaris* genome assembly and genome annotation will provide a valuable resource to the cephalopod and neuroscience communities.

Here we describe a chromosome-scale genome assembly and annotation of the common octopus, *O. vulgaris*. We have validated our assembly using available chromosome-scale genomes of octopus species (Li et al. 2020, Albertin et al. 2022; Jiang et al. 2022). Our analyses reveal large-scale chromosomal homologies, yet a pattern of local rearrangement within chromosomes between species.

## Materials and methods

### Sample Collection

One adult male *Octopus vulgaris* (780 g body weight, specimen tube3-27.05.21-GP, BioSamples ERS14895525 and ERS14895526) was collected in the Gulf of Naples, Italy (40°48’04.1”N 14°12’32.7”E) by fishermen in May 2021. The animal was immediately sacrificed humanely following EU guidelines and protocols for collection of tissues from wild animals (Andrews et al. 2013; Fiorito et al. 2015) (see Data Availability for animal welfare information). The central brain masses (optic lobes, OL; supra-, SEM; sub-esophageal, SUB) were dissected out (ERS14895525), and the spermatophores (ERS14895526) were collected as described in Zarrella et al. (2019). All dissections were carried out on a bed of ice in seawater, and the excised tissues were then weighed and flash-frozen in liquid nitrogen.

### High Molecular Weight Genomic DNA Extraction

High molecular weight genomic DNA (HMW gDNA) was extracted from frozen spermatophore sample (160 mg) (ERS14895526) using a salt-extraction protocol at the Stazione Zoologica Anton Dohrn (Italy) following Albertin et al. 2022. Briefly, two cryopreserved sample aliquots were each lysed for 3 hours at 55°C in separate tubes of 3 mL lysis buffer containing proteinase K. Then 1 mL of NaCl (5M) was added to each tube. The tubes were mixed by inversion then spun down for 15 minutes at 10,000 rcf. The supernatants were then transferred to a new tube and 2 volumes of cold ethanol (100%) was added. The DNA precipitate was then spooled, washed, resuspended in elution buffer (10 mM Tris, 0.1 mM EDTA, pH 8.5), and stored at 4°C. The DNA concentration was quantified using a Qubit DNA BR Assay kit (Thermo Fisher Scientific), and the purity was evaluated using Nanodrop 2000 (Thermo Fisher Scientific) UV/Vis measurements.

### 10X Genomics Library Preparation and Sequencing

A 10 ng aliquot of the spermatophore HMW DNA was used to prepare a 10X Genomics Chromium library (Weisenfeld et al. 2017) at the National Center for Genomic Analysis (Centre Nacional d’Anàlisi Genòmica - CNAG, Spain) using the Chromium Controller instrument (10X Genomics) and Genome Reagent Kits v2 (10X Genomics) following the manufacturer’s protocol. The library was indexed with both P5 and P7 indexing adaptors. The resulting sequencing library was checked that the insert size matched the protocol specifications on an Agilent 2100 BioAnalyzer with the DNA 7500 assay (Agilent).

The library was sequenced at CNAG with an Illumina NovaSeq 6000 with a read length of 2×151bp, and was demultiplexed with dual indices (Supplementary Data 1).

### Long-read Whole Genome Library Preparation and Sequencing

The spermatophore HMW DNA was also used to prepare one Oxford Nanopore Technologies (ONT) 1D sequencing library (kit SQK-LSK110) at CNAG. Briefly, 2.0 μg of the HMW DNA was treated with the NEBNext FFPE DNA Repair Mix (NEB) and the NEBNext UltraII End Repair/dA-Tailing Module (NEB). ONT sequencing adaptors were then ligated to the DNA, then the DNA was purified with 0.4X AMPure XP Beads and eluted in Elution Buffer.

Two sequencing runs were performed at CNAG on an ONT PromethIon 24 using ONT R9.4.1 FLO-PRO 002 flow cells. The libraries were sequenced for 110 hours. The quality parameters of the sequencing runs were monitored by the MinKNOW platform version 21.05.8 (Oxford Nanopore Technologies) and base called with Guppy, version 5.0.11 (available through https://community.nanoporetech.com) (Supplementary Data 1).

### Omni-C Library Preparation and Sequencing

A DoveTail Genomics Omni-C library was prepared at SciLifeLab (Solna, Sweden) using the flash-frozen brain tissue from the same individual used to generate the ONT long reads and 10X Genomics chromium reads (ERS14895525). One hundred milligrams of brain tissue were pulverized to a fine powder using a mortar and pestle under liquid nitrogen. Two 20 mg aliquots of the pulverized tissue were fixed in PBS with formaldehyde and disuccinimidyl glutarate (DSG), and were prepared according to the manufacturer’s protocol as two separate libraries. To increase the final complexity, the two libraries bound to streptavidin beads were pooled together into a single tube prior to P7 indexing PCR. The amplified library was sequenced at SciLifeLab on an Illumina NovaSeq 6000 with a read length of 2×150 bp, and was demultiplexed with one index (Supplementary Data 1).

### Nuclear Genome Assembly

Sequencing produced 77Gb of ONT WGS reads (27.5x coverage) and 230.25 Gb of 10X Genomics linked reads (77.7x coverage). These data were assembled with the CNAG Snakemake assembly pipeline v1.0 (https://github.com/cnag-aat/assembly_pipeline) to obtain an optimal base assembly for further Hi-C scaffolding. In brief, this pipeline first preprocessed the 10X reads with *LongRanger basic* v2.2.2 (https://github.com/10XGenomics/longranger) and filtered the ONT reads with *FiltLong* v0.2.0 (https://github.com/rrwick/Filtlong), and then the ONT reads were assembled with both *Flye* v2.9 (Kolmogorov et al. 2019) and *NextDenovo* v2.4.0 (Hu et al. 2023). The following evaluations were run on both assemblies and after each subsequent step of the pipeline: BUSCO v5.2.2 (Manni et al. 2021) with *metazoan_odb10* and *Merqury* v1.1 (Rhie et al. 2020) to estimate the consensus accuracy (QV) and k-mer statistics, and *fasta-stats.py* for contiguity statistics. The best contig assembly was obtained with *NextDenovo* (see assembly metrics Supplementary Data 2), so the remaining steps of the pipeline were run on this assembly (Supplementary Figure 1 and Supplementary Data 2).

The assembly was polished with 10X Illumina and ONT reads using *Hypo* v1.0.3 (Kundu et al. 2019); collapsed with *purge_dups* v1.2.5 (Guan et al. 2020); then scaffolded with the 10X chromium reads using *Tigmint* v1.2.4 (Jackman et al. 2018), *ARKS* v1.2.2 (Coombe et al. 2018) and *LINKS* v1.8.6 (Warren et al. 2015) following the Faircloth’s Lab protocol (http://protocols.faircloth-lab.org/en/latest/protocols-computer/assembly/assembly-scaffolding-with-arks-and-links.html). The specific parameters and versions used to assemble the *O. vulgaris* specimen are listed in Supplementary Data 3. Finally, 310 scaffolds shorter than 1 Kb were removed from the assembly. This assembly was used for scaffolding with Omni-C data.

### Omni-C Scaffolding

The Omni-C reads (863.85 million read pairs) were then mapped to the assembly (Supplementary Data 4) using the recommended procedure from Dovetail Genomics (https://omni-c.readthedocs.io/en/latest/fastq_to_bam.html). In short, the reads were mapped to the reference using *bwa mem* v0.7.17-r1188 (Li 2013) with flags *-5SP-T0*, converted to a sorted .*bam* file, and filtered to reads with a minimum mapping quality of 30 with *samtools* v1.9 (Li et al. 2009) with *htslib* v1.9, and filtered to keep uniquely mapping pairs with *pairtools* v0.3.0 (Open2C et al. 2023). The minimum mapping quality threshold of 30 was used to accommodate for the organism’s heterozygosity and repetitiveness (1.22% and 68.68%, respectively. see supplementary table Supplementary Data 5). After excluding PCR duplicates and improperly mated reads with *pairtools*, 231.59 million Hi-C read pairs were used to scaffold the assembly with *YaHS* v1.1 (Zhou et al. 2023) in the default mode, thus initially detects and corrects errors in contigs, introducing breaks at misjoins.

### Generation of the Hi-C Heatmaps and Manual Curation

We then manually curated the scaffolded assembly using an editable Hi-C heatmap to improve the assembly’s quality and to correct misassemblies. The process described below was repeated for five rounds until there were no obvious improvements to make based on the Hi-C heatmap signal.

*Chromap* v0.2.3 (Zhang et al. 2021) was used to align the Omni-C reads to the genome with a read alignment quality cutoff of Q0. The resulting *.pairs* file (quality cutoffs: 2,10) was converted using *awk* v 4.2.1(Aho et al. 1988) to a *.longp* file, a format used by *Juicebox Assembly Tools* (Dudchenko et al. 2018). We ran the script *run-assembly-visualizer.sh* from the *3D-DNA* pipeline (Dudchenko et al. 2017) on the *.longp* file to generate a *.hic* file. The *generate-assembly-file-from-fasta.awk* script from the *3D-DNA* pipeline (Dudchenko et al. 2017), and the *assembly-from-fasta.py* from the *Artisanal* pipeline (Bredeson et al. 2022) were used to generate the *.assembly* files necessary to curate the *.hic* heatmap file in *Juicebox Assembly Tools* (Dudchenko et al. 2018).

The resulting *.hic* heatmap file was visualized using the visualization tool *Juicebox* v1.11.08 (Durand et al. 2016). Using the signal in the Hi-C heatmap we corrected the order and orientation of contigs within the chromosome-scale scaffolds, and placed small contigs and scaffolds onto the chromosome-scale scaffolds. A new *.fasta* assembly was generated from the corrected *.assembly* file by using the *assembly-to-fasta.py* script from the *Artisanal* pipeline.

The corrected assembly was aligned to the chromosome-scale *O. sinensis* (GCA_006345805.1) (Li et al. 2020), *O. bimaculoides* (GCA_001194135.2) (Albertin et al. 2022), and *A. fangsiao* (Jiang et al. 2022) genomes using *minimap2* v2.24 (Li, 2018), *snakemake* v7.19.1-3.11.1 (Köster and Rahmann 2012) and the *snakemake* script *GAP_dgenies_prep* (https://doi.org/10.5281/zenodo.7826771). The resulting *.paf* file was visualized with *D-GENIES* v1.4.0 (Cabanettes and Klopp 2018). Regions of the *O. vulgaris* chromosome-scale scaffolds that had ambiguous Hi-C heatmap signal, or regions that had no obvious homology to other *Octopus* spp. chromosome-scale scaffolds were removed from the chromosome-scale scaffolds and retained as smaller scaffolds at the end of the genome assembly *.fasta* file. Scaffolds were renamed based on homology with *O. bimaculoides* chromosomes.

### Decontamination

After curation, we ran the *BlobToolKit* INSDC pipeline (Challis et al. 2020), using the NCBI *nt* database (updated on December 2022) and the following BUSCO *odb10* databases: eukaryota, fungi, bacteria, metazoa and mollusca. This analysis identified 226 scaffolds either matching the phylum Mollusca or having no-hit in the database (Supplementary Figure 2). A total of 47 small scaffolds matching other phyla (Supplementary Data 6 and Supplementary Figure 3) were considered contaminants and removed from the assembly. This scaffolded and decontaminated assembly was then carried forward for annotation and comparative analyses, and is available at https://denovo.cnag.cat/octopus and the INSDC (ENA, NCBI, and DDBJ) accession number GCA_951406725.1.

### Nuclear Genome Annotation

The gene annotation of the octopus genome assembly was obtained by combining transcript alignments, protein alignments, and *ab initio* gene predictions as described below. A flowchart of the annotation process is shown in Supplementary Figure 4.

Repeats present in the genome assembly were annotated with *RepeatMasker* v4-1-2 (Smit et al. 2013-2015) using the custom repeat library available for Mollusca. Moreover, a new repeat library specific to the assembly was made with *RepeatModeler* v1.0.11. After excluding repeats from the resulting library that were part of repetitive protein families by performing a *BLAST* (Altschul et al. 1990) search against *Uniprot*, *RepeatMasker* was rerun with this new library to annotate species-specific repeats.

PacBio Iso-Seq reads from several developmental stages were downloaded from NCBI (PRJNA718058, PRJNA791920, PRJNA547720) (García-Fernández et al. 2019; Deryckere et al. 2021; Zolotarov et al. 2022). Bulk RNA-seq from an adult octopus (Petrosino et al. 2022) was downloaded from the *ArrayExpress* database under accession number E-MTAB-3957. The short and long reads were aligned to the genome using *STAR* v-2.7.2a (Dobin et al. 2013) and *minimap2* v2.14 (Li, 2018) with the option *-x splice:hq*. Transcript models were subsequently generated using *Stringtie* v2.1.4 (Pertea et al. 2015) on each .*bam* file, and then all the transcript models were combined using *TACO* v0.6.3 (Niknafs et al. 2017). High-quality junctions to be used during the annotation process were obtained by running *Portcullis* v1.2.0 (Mapleson et al. 2018) after mapping with *STAR* and *minimap2*. Finally, *PASA* assemblies were produced with *PASA* v2.4.1 (Haas et al. 2008). The *TransDecoder* program, part of the *PASA* package, was run on the *PASA* assemblies to detect coding regions in the transcripts.

The complete proteomes of *O. vulgaris*, *O. bimaculoides,* and *Sepia pharaonis* were downloaded from *UniProt* in October 2022 and aligned to the genome using *Spaln* v2.4.03 (Iwata and Gotoh 2012).

*Ab initio* gene predictions were performed on the repeat-masked assembly with two different programs: *Augustus* v3.3.4 (Stanke et al. 2006) and *Genemark-ES* v2.3e (Lomsadze et al. 2014) with and without incorporating evidence from the RNA-seq data. Before gene prediction, *Augustus* was trained with octopus-specific evidence. The gene candidates used as evidence for training *Augustus* were obtained after selecting *Transdecoder* annotations that were considered complete and did not overlap repeats, clustering them into genes, and selecting only one isoform per gene. These candidates were aligned to the *Swissprot* NCBI database with *blastp* v2.7.1 (Altschul et al. 1990) to select only those with homology to proteins. The final list of candidate genes was made of 1764 genes with *BLAST* hits to known proteins with e-values smaller than 10^-9^ and greater than 55% identity.

Finally, all the data were combined into consensus CDS models using *EvidenceModeler* v1.1.1 (EVM) (Haas et al. 2008). Additionally, UTRs and alternative splicing forms were annotated via two rounds of *PASA* annotation updates. Functional annotation was performed on the annotated proteins with *Blast2go* v1.3.3 (Conesa et al. 2005). First, a *DIAMOND* v2.0.9 *blastp* (Buchfink et al. 2021) search was made against the *nr* database. Furthermore, *Interproscan* v5.21-60.0 (Jones et al. 2014) was run to detect protein domains on the annotated proteins. All these data were combined by *Blast2go* v1.3.3, which produced the final functional annotation results.

Identification of long non-coding RNAs (lncRNAs) was done by first filtering the set of *PASA*-assemblies that had not been included in the annotation of protein-coding genes to retain those longer than 200bp and not covered more than 80% by repeats. The resulting transcripts were clustered into genes using shared splice sites or significant sequence overlap as criteria for designation as the same gene.

### Nuclear Genome and Annotation Completeness Assessment

The final *O. vulgaris* genome assembly, the annotated transcripts, the proteins from the annotated transcripts, and the other available octopus genomes were assessed for completeness using BUSCO databases as described above (Materials and Methods - Genome Assembly). To compare the qualities of each assembly, we used *fasta_stats* (Chapman et al. 2011) shown in (Table 1). We calculated the percentage of bases in the chromosome-scale scaffolds (Table 1) with *bioawk* v1.0 (https://github.com/lh3/bioawk).

**Table 1.** Octopus genome assembly statistics.

| Assembly | Number of scaffolds | Number of contigs | Scaffold sequence total | Scaffold N50/L50 | Number of scaffolds > 50 KB | % of the bases in chromosome-scaled scaffolds |
| --- | --- | --- | --- | --- | --- | --- |
| Final chromosome-scale *O. vulgaris* genome | 226 | 2758 | 2800.4 MB | 118.9 MB/9 | 57 | 99.34 |
| Pre-curation scaffolded assembly *O. vulgaris* | 768 | 2776 | 2801.6 MB | 118.3 MB/9 | 296 | 95.82 |
| Chromosome-scale *O. bimaculoides* (Albertin et al. 2022) | 145326 | 713915 | 2342.5 MB | 96.9 MB/9 | 85 | 95.46 |
| Chromosome-scale *O. sinensis* (Li et al., 2020) | 13516 | 20491 | 2719.2 MB | 105.9 MB/10 | 1800 | 86.09 |
| Chromosome-scale *A. fangsiao* (Jiang et al. 2022) | 6409 | 9099 | 4341.1 MB | 169.7 MB/10 | 1769 | 93.05 |

### Mitogenome Assembly and Annotation

To assemble the mitochondrial genome we employed a strategy that uses a reference bait to select the mitochondrial nanopore reads, assembles those reads into a single circular contig, and then performs two rounds of polishing. To obtain the mitochondrial sequences, all ONT reads with a mean quality of ≥10 were mapped with *minimap2* v2.24 (Li 2018) against the circular complete, 15,744 bp mitochondrial genome of another specimen of *O. vulgaris* (NC_006353.1) (Yokobori 2004) with the minimap2 parameter *-ax map-ont*. We retained all reads with a mapping quality ≥ 13. Approximately 5,000 ONT reads passed these filters including 15 reads accounting for 181,644 total basepairs (12x coverage) with a mean length of 12,112 bp.

All the retained ONT reads were assembled with *Flye* v2.9 (Kolmogorov et al. 2019) using the options *flye --scaffold -i 2 -g 15744 --nano-raw --min-overlap 7000*. This produced one circular contig. The -*i 2* option specified for flye caused this contig to be polished twice with the input ONT reads. After polishing the length of the circular contig was 15,651 bp, and a web blastn search revealed that it spanned the length of the NC_006353.1 mitochondrial genome. The circular mitogenome contig was rotated and oriented as follows. First, we annotated the contig using *MITOS* v2.1.3 (Bernt et al. 2013) with parameters *-c 5 --linear --best -r refseq81m*. Second, we use the coordinates in the *results.bed* file to orient the mitogenome, so it starts with the conventional tRNA Phenyl-Alanine (trnF) (Formenti et al. 2021).

To evaluate the assembly accuracy, we first aligned the selected ONT reads back to the assembly with *minimap2* and visually inspected the alignment with *IGV* v2.14.1 (Robinson et al. 2023). Finally, the xcOctVulg1 mitogenome was aligned against the mitogenome of other species using *DNAdiff* v1.3 from *mummer* package v3.23 (Kurtz et al. 2004). These species included the mitogenomes of another specimen of *O. vulgaris* (NC_006353.1), *O. sinensis* (NC_052881.1), *O. bimaculoides* (NC_029723.1), and *A. fangsiao* (AB240156.1). From these pairwise alignments, we calculated the percent identity.

## Results and Discussion

### DNA Sequencing

Sequencing the ONT WGS library yielded 8.3 million ONT PromethIon reads containing 82.57 billion base pairs (Gbp) with 29.47x coverage per library. Sequencing of the 10X Genomics Chromium library yielded 762 million read pairs containing 228.69 Gbp with 81.64x coverage per library. The Omni-C library sequencing yielded 863.85 million read pairs, containing 259.16 Gbp of data with 33.02X coverage. Details about sequence data can be found in Supplementary Data 1.

### Manual Curation and Decontamination of the Assembly

Manually curating the genome assembly improved the quality of the final assembly, as 495 scaffolds were placed in the chromosome-scale scaffolds, and 47 additional scaffolds were removed through the contamination analysis (Table 1). The final 2.80 Gb assembly, xcOctVulg1.1, has a scaffold N50 of 118.9 Mb, an N90 of 18.2 Mb, QV39 and gene completeness estimated using BUSCO v5.3.2 with *mollusca_odb10* of C:86.5% [S:85.8%, D:0.7%], F:3.4%, M:10.1%, n:5295 (Fig. 1C). The BUSCO score with *metazoa_odb10* for the final assembly is C:92.3% [S:91.8%, D:0.5%], F:2.7%, M:5.0% (Table 2). The statistics for all intermediate assemblies are shown in Supplementary Data 2. Also, in Supplementary Figure 3 we show that the final assembly has been properly decontaminated.

**Figure 1.**
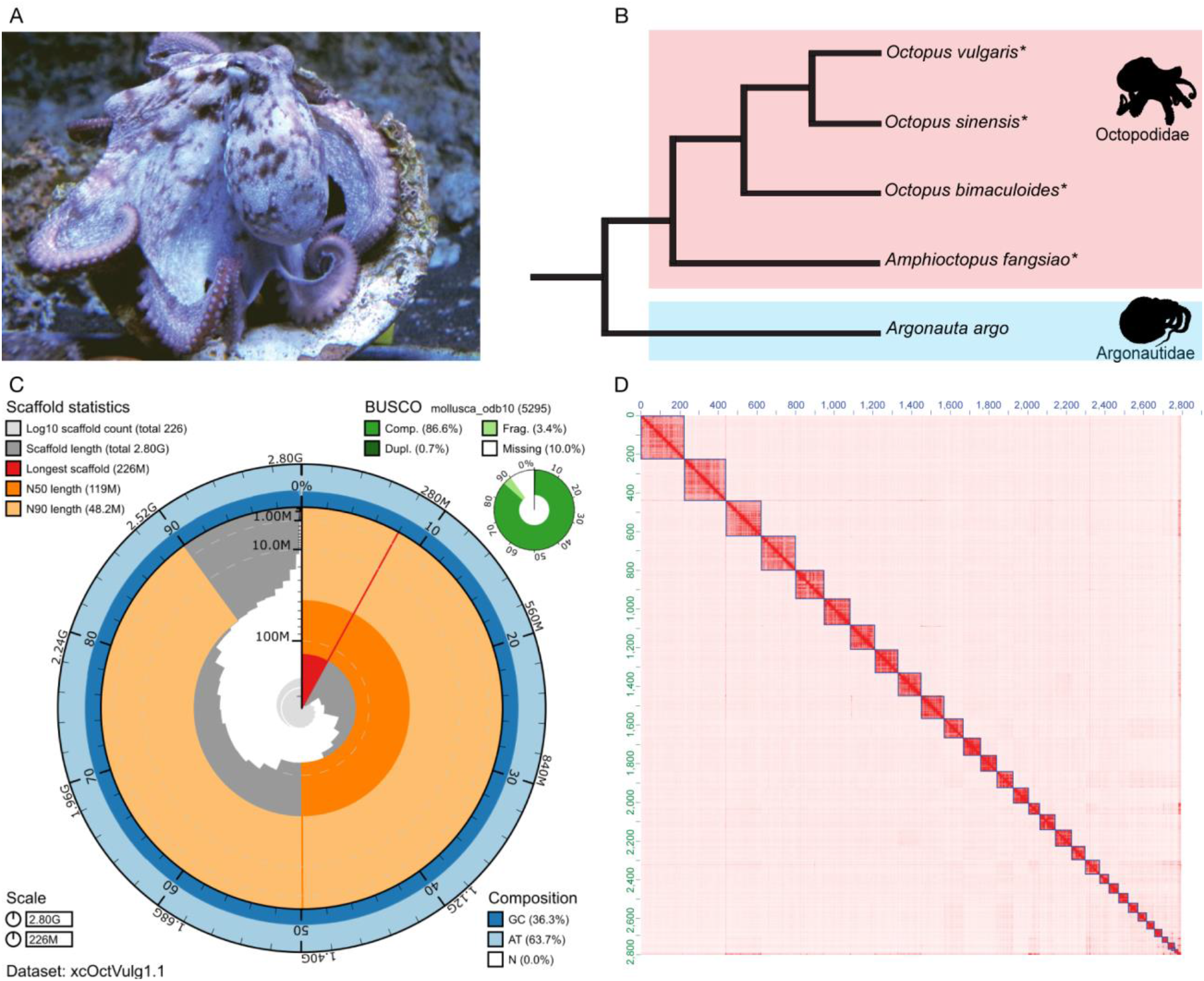
*Octopus vulgaris* assembly statistics and quality control. (A) A specimen of *O. vulgaris*. (B) A cladogram showing the phylogenetic relationship between the compared species and the family Argonautidae as an outgroup (Taite et al. 2023). Chromosome-scale genome assemblies are available for the starred species (*). (C) The snail plot shows that the final version of the chromosome-scale *O. vulgaris* assembly has N50 of 119Mb, the longest scaffold is 225Mb long, and a BUSCO score for complete genes of 86.6% against the *mollusca_odb10* database. (D) The Hi-C heatmap of the final genome assembly shows 30 chromosome-scale scaffolds with very few sequences in unplaced scaffolds. Photography credit © Antonio, Valerio Cirillo (BEOM SZN), 2023 (A).

**Table 2.** Octopus *metazoa_odb10* BUSCO scores.

| Genome | Complete BUSCO | Single BUSCO | Duplicated BUSCO | Fragmented BUSCO | Missing BUSCO |
| --- | --- | --- | --- | --- | --- |
| Chromosome-scale *O. vulgaris* | 92.3%<br>[881] | 91.8%<br>[876] | 0.5%<br>[5] | 2.7%<br>[25] | 5.0%<br>[48] |
| Contig-level *O. vulgaris*<br>(Zarrella et al. 2019) | 63.1%<br>[602] | 62.6%<br>[597] | 0.5%<br>[5] | 24.8%<br>[237] | 12.1%<br>[115] |
| Chromosome-scale *O. bimaculoides*<br>(Albertin et al. 2022) | 94.6%<br>[903] | 94.2%<br>[899] | 0.4%<br>[4] | 3.2%<br>[31] | 2.2%<br>[20] |
| Chromosome-scale *O. sinensis* (Li et al., 2020) | 95.7%<br>[913] | 90.5%<br>[863] | 5.2%<br>[50] | 2.6%<br>[25] | 1.7%<br>[16] |
| Chromosome-scale *A. fangsiao*<br>(Jiang et al. 2022) | 93.5%<br>[892] | 91.6%<br>[874] | 1.9%<br>[18] | 3.5%<br>[33] | 3.0%<br>[29] |

### The Octopus Karyotype

The genome assembly from this study contains 30 large scaffolds with Hi-C heatmap signal (Fig. 1D) that is consistent with each scaffold representing a single chromosome and resembles the Hi-C heatmaps of other chromosome-scale octopus genome assemblies (Li et al. 2020; Albertin et al. 2022; Jiang et al. 2022). The first reported *O. vulgaris* karyotypes from Japan and Italy were 1n=28 chromosomes (Inaba 1959; Vitturi et al. 1982), but later studies also using *O. vulgaris* individuals sampled in Japan reported at 1n=30 (Gao and Natsukari 1990). The karyotype 1n=30 has been reported in four other octopus species: *Callistoctopus minor, Amphioctopus fangsiao, Cistopus sinensis*, and *Amphioctopus areolatus* (Gao and Natsukari 1990; Adachi et al. 2014; Wang and Zheng 2017). The only exception is *Hapalochlaena maculosa* which does not have a confirmed karyotype, but 47 linkage groups were suggested for this species (Whitelaw et al. 2022).

In light of the recent taxonomic designation of a new species *O. sinensis* (East Asian Common Octopus) from the previously synonymous *O. vulgaris* (Gleadall 2016; Amor et al. 2017, 2019; Amor 2023), this suggests that the reported *O. vulgaris* karyotypes probably belong to *O. sinensis*. Dot plot analyses, described below, show that *O. vulgaris* and *O. sinensis* share 30 homologous, largely collinear, chromosomes (Fig. 2).

**Figure 2.**
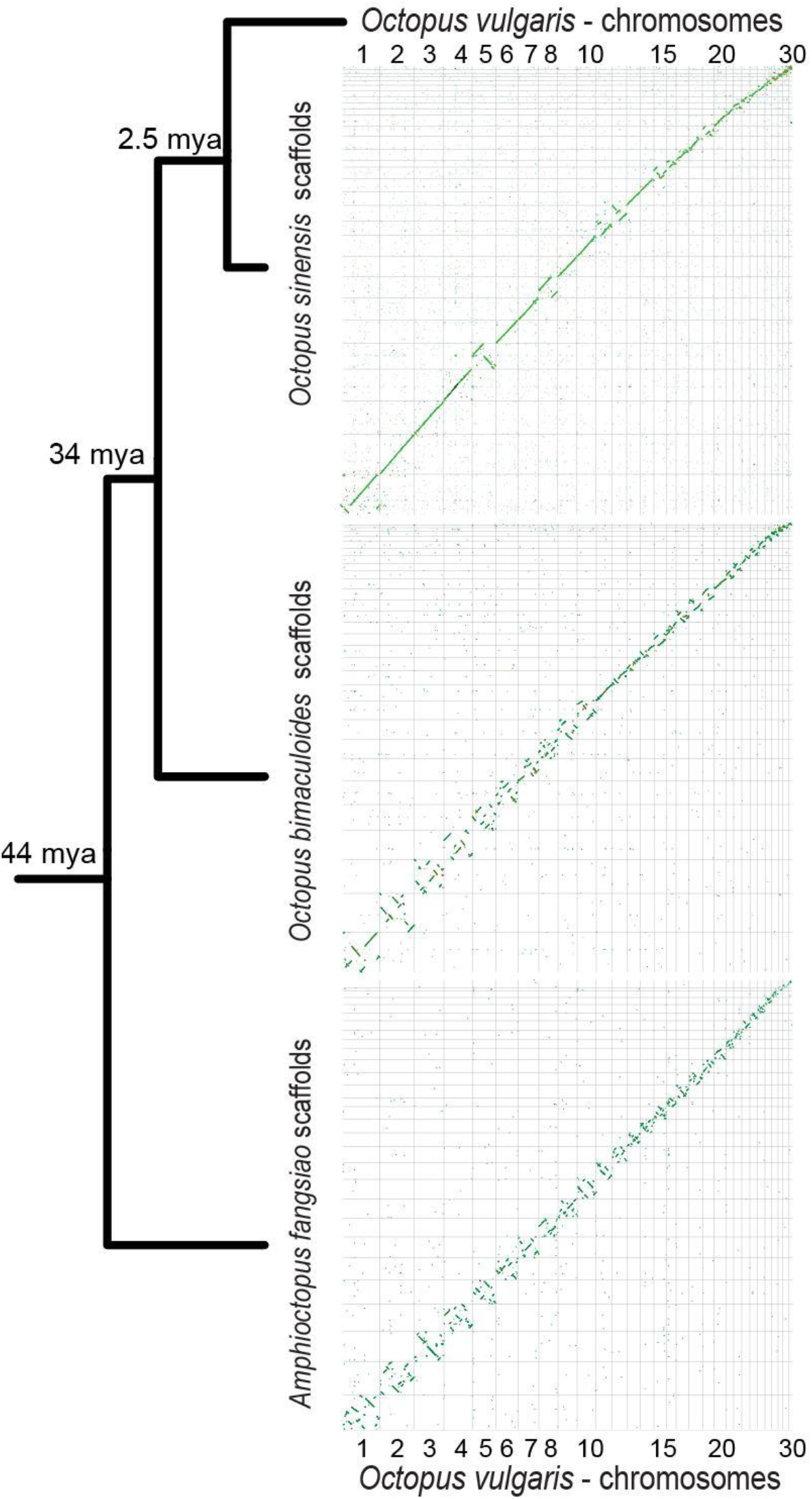
Comparative analyses of available chromosome-scale Octopodidae genomes. The figure shows the inferred phylogenetic relationship (Amor et al. 2017; Jiang et al. 2022; Taite et al. 2023) and the inferred divergence times (Amor et al. 2019; Jiang et al. 2022) of four octopus species. The diagrams show genome-genome alignments for each species compared to *O. vulgaris*.

The final version of the *O. vulgaris* genome was aligned to the genomes of three octopus species, *O. sinensis*, *O. bimaculoides*, and *A. fangsiao* (Fig. 2). *O. vulgaris* and *O. sinensis* have a less diverged genome sequence and few inversions between homologous, collinear chromosomes. General chromosomal collinearity was also observed in comparison to *O. bimaculoides* (Fig. 2). We have found large-scale inversions (megabase-scaled, larger than 1Mb) throughout the genomes of two species. The overall sequence similarity is lower compared to the previous pair, and a greater number of chromosomal rearrangements are present, confirming that they are more diverged. This is expected considering that *O. bimaculoides* and the *O. vulgaris*-*O. sinensis clade* diverged around 34 million years ago (mya) (Jiang et al. 2022), while *O. sinensis* and *O. vulgaris* diverged just 2.5 mya (Amor et al. 2019). In Figure 2, the collinearity between *O. vulgaris* and *A. fangsiao* chromosomes is visible. As expected, as *A. fangsiao* is the most distant to *O. vulgaris* of the compared species, the genomes are even more rearranged.

Our whole-genome alignment analyses support the hypothesis that *O. vulgaris, O. sinensis*, *O. bimaculoides,* and *A. fangsiao* share 30 homologous chromosomes (Fig. 2). Given the divergence time of these species, these results suggest that the karyotype of the common ancestor of this clade, and perhaps the common ancestor of octopuses, also had 30 chromosomes that still exist in extant species.

Karyotype stability was described in the squid lineage (Decapodiformes) on loliginid and sepiolid squids (Albertin et al. 2022). This study has suggested that the smaller karyotype found in octopuses (1n=30) compared to squids (1n=46) results from secondary fusions of a more ancestral squid chromosomal complement. Recently, it has been suggested that chromosomal fusions impact recombination, as well as chromosomal nuclear occupancy, in mice (Vara et al. 2021). Therefore, chromosomal fusions in the common ancestor of the octopus lineage might be one of the drivers of diversification, as this changes the chromosomal interactions and is hypothesized to lead to the formation of novel regulatory units (Vara et al. 2021). Such events are important in light of understanding the emergence of octopus-specific traits. We infer from the genome-genome comparisons that a similar pattern of intrachromosomal rearrangements with the conservation of individual chromosomes is seen in octopus species, as described in squids (Albertin et al. 2022). However, the loliginids and sepiolids are estimated to have diverged 100 mya (Albertin et al. 2022), while the genera *Octopus* and *Amphioctopus* are estimated to have diverged 44 mya (Jiang et al. 2022). Therefore, a more-distant species chromosome-scale genome is needed to claim karyotype stasis in Octopodiformes. Nevertheless, future comparative studies of the genomes of these closely-related species will shed light on the evolutionary history of octopuses as a separate lineage of coleoid cephalopods. In addition to this, *O. vulgaris* is a model animal in neurobiological studies, and having a high-quality genome will facilitate further studies of the cephalopod brain.

### Nuclear Genome Annotation

In total, we annotated 23,423 protein-coding genes that produce 31,799 transcripts (1.36 transcripts per gene) and encode 30,121 unique protein products. We were able to assign functional labels to 53.5% of the annotated proteins. The annotated transcripts contain 8.42 exons on average, with 87% of them being multi-exonic (Table 3). In addition, 1,849 long non-coding transcripts have been annotated. The number of protein-coding genes annotated here is slightly lower than those reported for other octopus genome assemblies, like *O. sinensis* (Li et al. 2020). After checking the general statistics of both annotations (Table 3), we can observe that the genes annotated here tend to be longer (both in the number of exons and global length). After comparing both methods, the main difference that we believe is responsible for this difference in length is the source of the transcriptomic data, the inclusion of long-read Iso-seq data in the annotation process is known to result in less fragmented and longer annotations.

**Table 3.** Genome annotation statistics.

|  | **OctVul6B annotation** | **Osinensis**<br>**ASM634580v1**<br>**(Li et al. 2020)** |
| --- | --- | --- |
| **Number of protein-coding genes** | 23,423 | 31,676 |
| **Median gene length (bp)** | 20,288 | 4,403 |
| **Number of transcripts** | 31,799 | 31,676 |
| **Number of exons** | 168,570 | 184,658 |
| **Number of coding exons** | 161,430 | 180,943 |
| **Median UTR length (bp)** | 1,255 | 441 |
| **Median intron length (bp)** | 2,467 | 1,520 |
| **Exons/transcript** | 8.42 | 5.83 |
| **Transcripts/gene** | 1.36 | 1 |
| **Multi-exonic transcripts** | 87% | 81% |
| **Gene density (gene/Mb)** | 8.36 | 11.7 |

### Nuclear Genome and Annotation Completeness Assessment

The BUSCO score was calculated for the *O. vulgaris*, *O. bimaculoides*, *O. sinensis*, and *A. fangsiao* genomes. For the chromosome-scale *O. vulgaris* genome, the BUSCO score for a whole-genome nucleotide sequence using the metazoan reference dataset was 92.3% for complete genes (954 core genes). The full score can be seen in Table 2. This is an improvement considering the BUSCO score of the previous *O. vulgaris* genome assembly (GCA_003957725.1) for complete genes was 63.1% (Zarrella et al. 2019). Additionally, we assessed the completeness of the annotated proteome and transcriptome by calculating the BUSCO score against the *metazoa_odb10* and *mollusca_odb10* databases (Supplementary Data 2).

### Mitogenome Assembly and Annotation

The mitogenome assembly of the *O. vulgaris* specimen (xcOctVulg1) has a length of 15,651 bp and contains 13 protein-coding, 23 ncRNA, 2 rRNA, and 21 tRNA genes. The ONT read alignment to the mitogenome shows a high consensus support for each nucleotide except for 16 positions (Supplementary Figure 5). These 16 positions are single nucleotide polymorphisms, not indels, and the base at each position is the base with the highest coverage in the reads at that position (Supplementary Figure 6). Therefore, the mitochondrial genome has a high per-base accuracy.

The percentages of identity (See Supplementary Data 7) between the *O. vulgaris* and other octopus mitochondrial genome sequences are consistent with the phylogeny topology (Fig. 2, Supplementary Data 7), and previous research on octopus taxonomy. The mitochondrial genome of the specimen collected in Japan and identified as *O. vulgaris* (NC_006353.1) shows a higher identity to *O. sinensis* (99.85%) than to our *O. vulgaris* specimen (96.79%). The 3.21% difference between the mitogenomes of the specimen from this study and NC_006353.1 is close to the estimated divergence rate (~2% divergence/million years (Arbogast and Slowinski 1998)) for *O. vulgaris* and *O. sinensis* (estimated time of divergence: 2.5mya (Amor et al. 2019). These results suggest that the specimen collected in Japan and identified as *O. vulgaris* (NC_006353.1) is more likely to be *O. sinensis*. This possibility is consistent with recent morphological, molecular, and geographic delimitations made between the *O. sinensis* and *O. vulgaris* species complex (Gleadall 2016; Amor et al. 2017, 2019; Amor 2023).

## Conclusion

*Octopus vulgaris* is an important emerging model in comparative neuroscience, cognition research, and evolutionary studies of cephalopods. The chromosome-scale genome assembly and annotation reported here provide an improved reference for single-cell multiomics and the study of non-coding regions and gene regulatory networks, that require the context of chromosome-scale sequences. This assembly and annotation will also facilitate many avenues of cephalopod research, in particular analyses of genome evolutionary trends in octopus and cephalopods within invertebrates. Furthermore, the chromosome-scale *O. vulgaris* genome assembly will allow the estimation of chromosome rearrangement rates, the emergence of novel coding and non-coding genes among octopuses, and the turnover rates of putative regulatory regions. The scientific interest in *O. vulgaris* as a model animal in many fields including (evolutionary) developmental biology and neuroscience will be facilitated by the availability of a high-quality genome.

These efforts may help bridge the traditional *O. vulgaris* research on neurobiology, behavior, and development to the molecular determinants involved in these fields.

## Data Availability Statement

The data are available at https://denovo.cnag.cat/octopus. On the INDSC databases (ENA, NCBI, DDBJ) the genome is available at accession GCA_951406725.1, and the data in BioProject PRJEB61268. Euthanizing cephalopods solely for tissue removal does not require authorization from the National Competent Authority under Directive 2010/63/EU and its transposition into National Legislation. Samples were taken from local fishermen, and humane killing followed principles detailed in Annex IV of Directive 2010/63/EU as described in the Guidelines on the Care and Welfare of Cephalopods (Fiorito et al. 2015). The sampling of octopuses from the wild included in this study was authorized by the Animal Welfare Body of Stazione Zoologica Anton Dohrn (Ethical Clearance: case 06/2020/ec AWB-SZN). Genomes of *O. sinensis* (GCA_006345805.1) (Li et al., 2020) and *O. bimaculoides* (GCA_001194135.2) (Albertin et al. 2022) were downloaded from NCBI, while the *A. fangsiao* genome (Jiang et al. 2022) was downloaded from Figshare (https://figshare.com/s/fa09f5dadcd966f020f3).

## Acknowledgments

The authors acknowledge that some of the computational results of this work have been achieved using the Life Science Compute Cluster (LiSC) of the University of Vienna. We also acknowledge support from the National Genomics Infrastructure in Stockholm funded by Science for Life Laboratory, the Knut and Alice Wallenberg Foundation and the Swedish Research Council, and NAISS/Uppsala Multidisciplinary Center for Advanced Computational Science for assistance with massively parallel sequencing and access to the UPPMAX computational infrastructure.

## Author Contributions

Dalila Destanovic = D.D.

Darrin T. Schultz = D.T.S.

Oleg Simakov = O.S.

Eve Seuntjens = E.S.

Ruth Styfhals = R.S.

Giovanna Ponte = G.P.

Tyler S. Alioto = T.S.A.

Fernando Cruz = F.C.

Jessica Gomez-Garrido = J.G.G.

Ivo Gut = I.G.

Marta Gut = M.G.

E.S., O.S. and G.P. conceived of the study design. D.D., D.T.S., O.S., T.S.A., M.G., and F.C. wrote the first draft of the manuscript. G.P. collected and dissected the octopus individual sequenced for this study. T.S.A., F.C., D.D., and D.T.S. assembled the genome. J.G.G. annotated the genome. D.D., J.G.G., and F.C. performed genomic analyses. D.D., D.T.S., F.C., J.G.G. and created the figures. All authors contributed to the interpretation, presentation, and writing of the manuscript.

## Supplementary Information

### This PDF file includes

Supplementary Figure 1 [Genome assembly pipeline]

Supplementary Figure 2 [Contamination analysis of the manually curated genome]

Supplementary Figure 3 [Contamination analysis of the final version of *O. vulgaris* genome]

Supplementary Figure 4 [Annotation workflow]

Supplementary Figure 5 [Alignment of ONT reads to the *O. vulgaris* mitochondrial genome.]

Supplementary Figure 6 [Coverage of ambiguous positions in the mitochondrial genome]

### Other Supplementary Data for this manuscript include the following

Supplementary Data 1 [Supplementary_Data_1.xlsx]

Supplementary Data 2 [Supplementary_Data_2.xlsx]

Supplementary Data 3 [Supplementary_Data_3.xlsx]

Supplementary Data 4 [Supplementary_Data_4.xlsx]

Supplementary Data 5 [Supplementary_Data_5.txt]

Supplementary Data 6 [Supplementary_Data_6.xlsx]

Supplementary Data 7 [Supplementary_Data_7.xlsx]

### Supplementary Figures

**Supplementary Figure 1.**
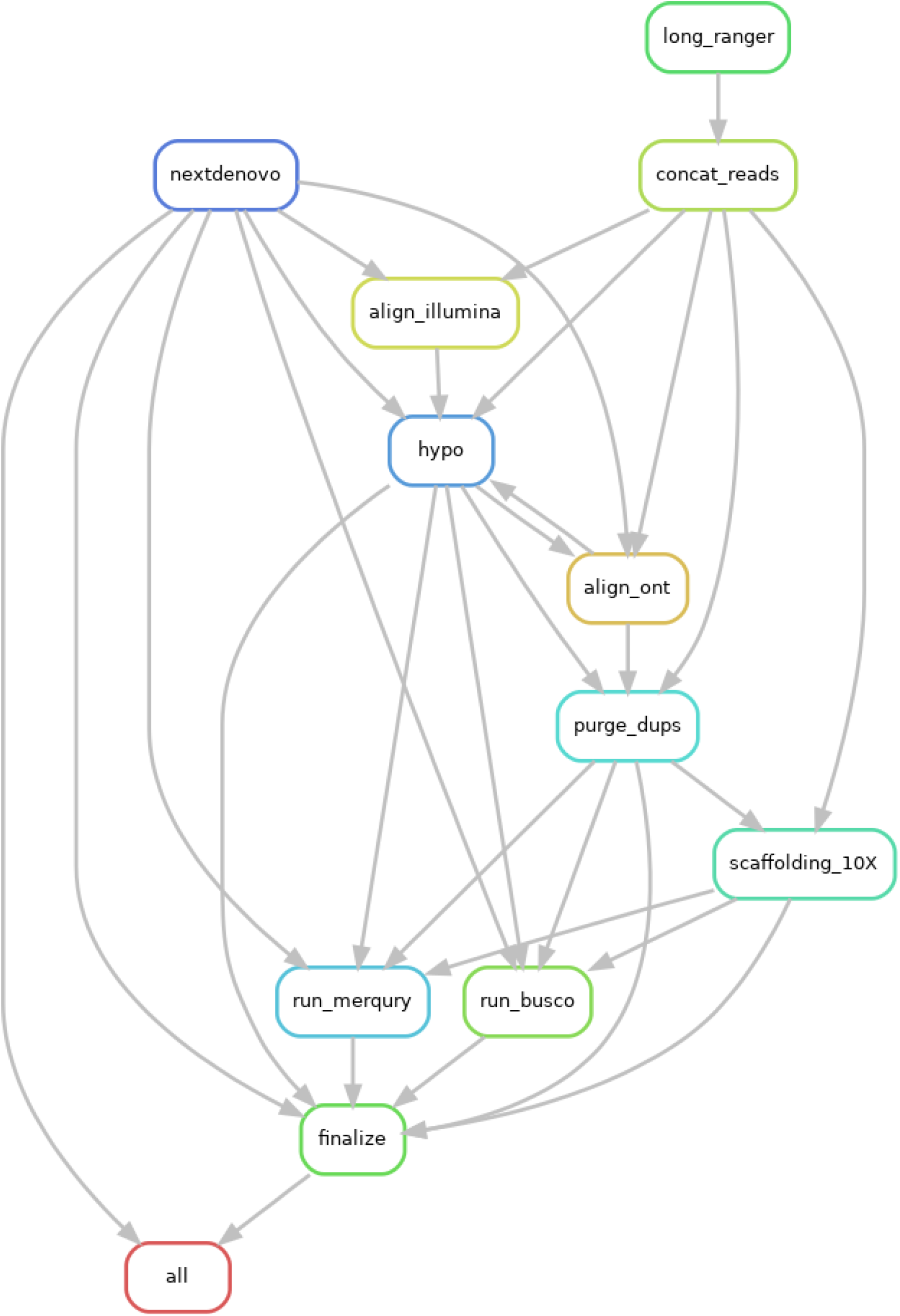
Genome assembly pipeline. Snakemake workflow is used to generate the scaffolded *Octopus genome* assembly.

**Supplementary Figure 2.**
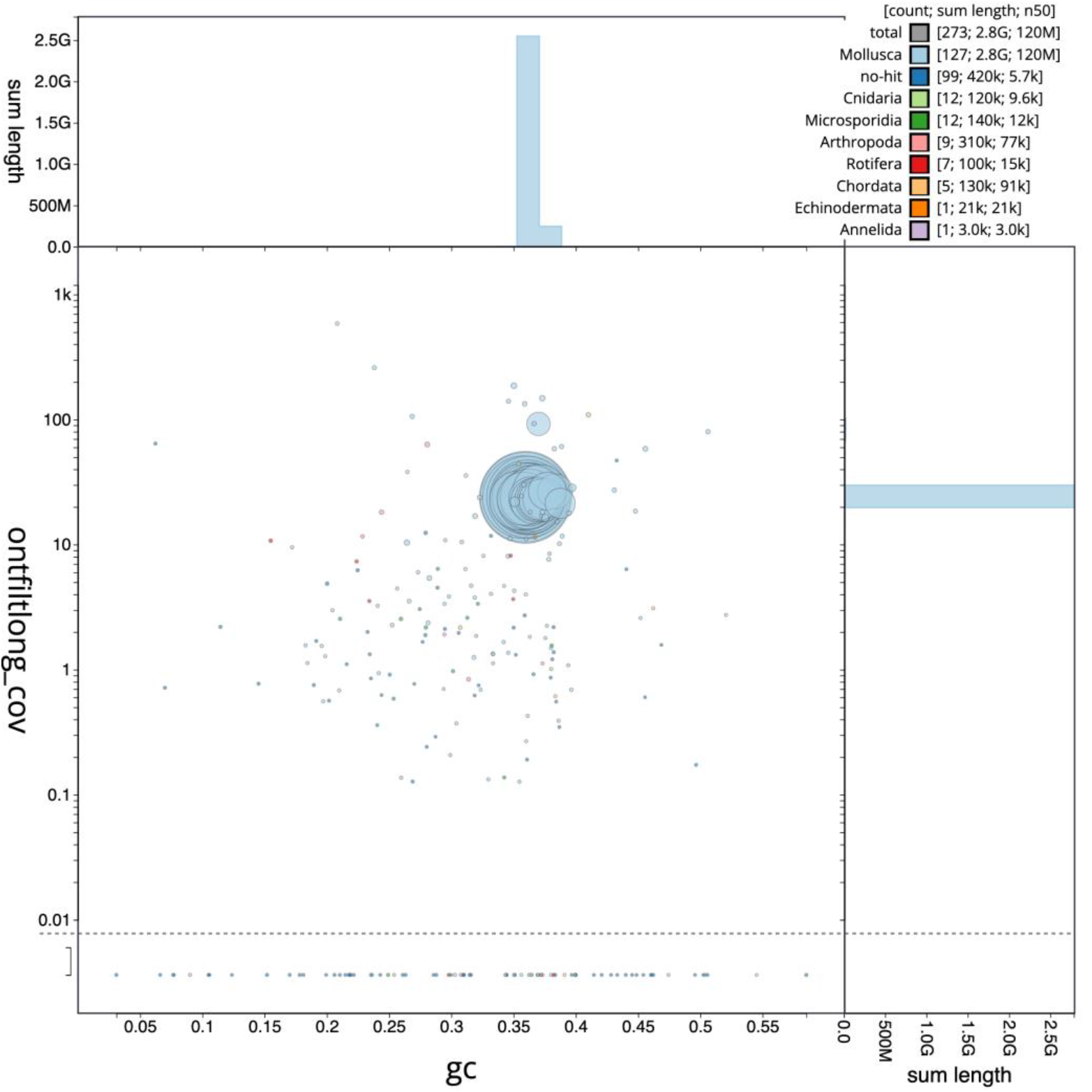
Contamination analysis of the manually curated genome. The original manually curated assembly had some sequences belonging to other phyla. These fragments were found in the unplaced scaffolds in the assembly.

**Supplementary Figure 3.**
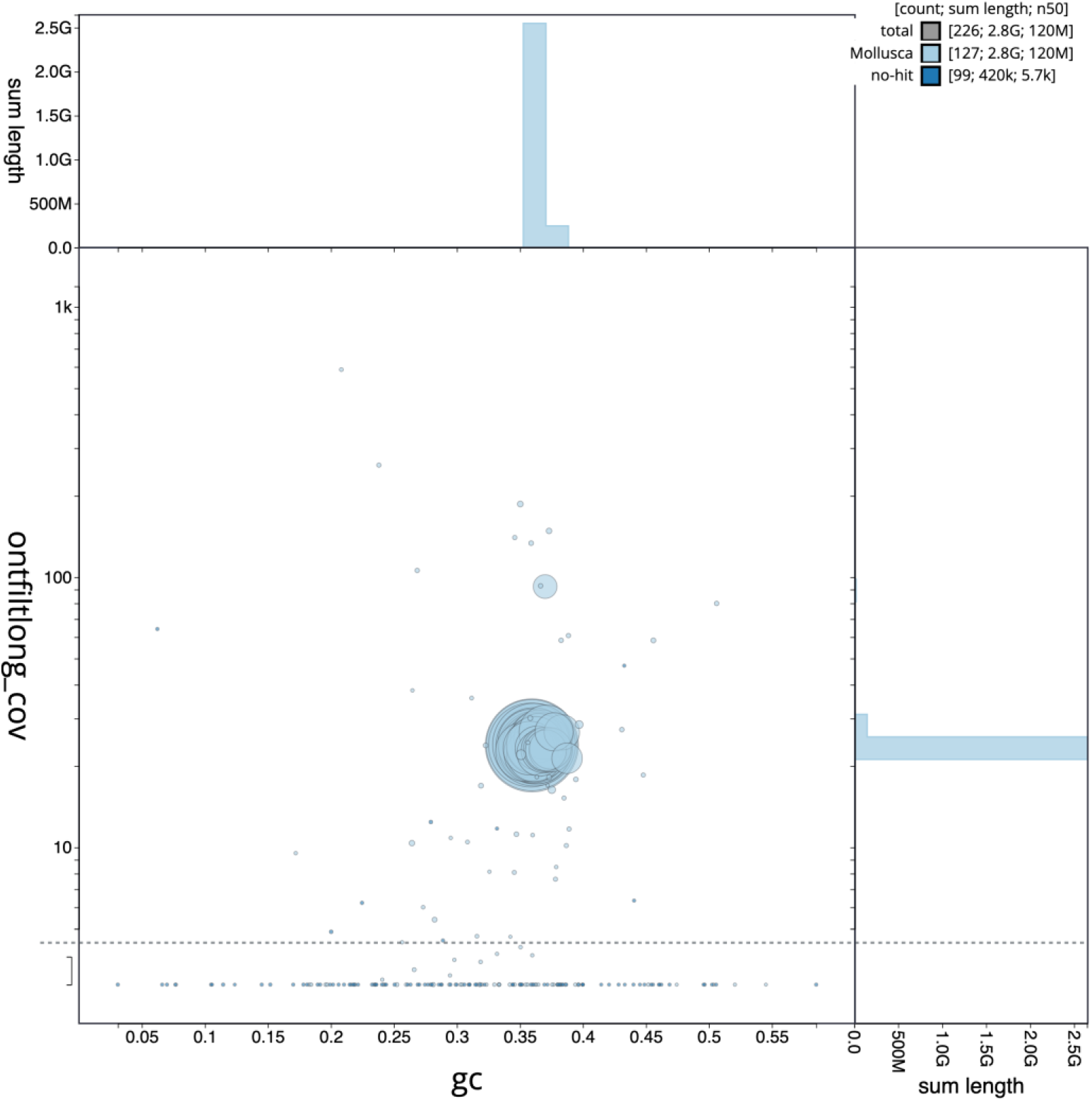
Contamination analysis of the final version of *O. vulgaris* genome. Decontamination of the assembly was successful, as the molluscan and no-hit sequences were kept. This is the final version of the chromosome-scale genome that was generated.

**Supplementary Figure 4.**
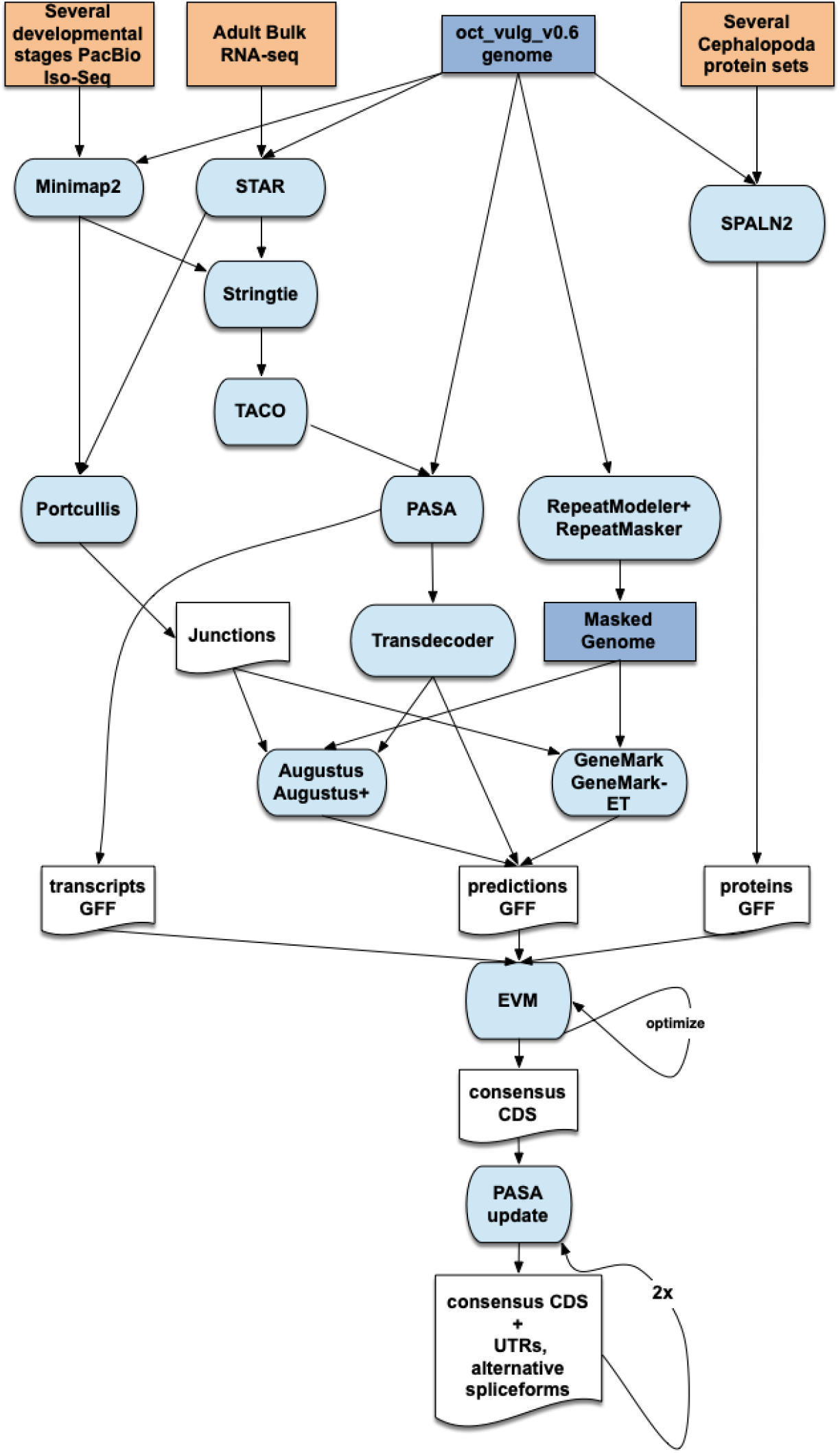
Annotation workflow. Combined data of protein sequences from different cephalopod species, and RNA sequences generated from adult and embryonic tissue were used to annotate the genome. This resulted in the annotation of CDS, UTRs, and alternative splice variants for the genome.

**Supplementary Figure 5.**
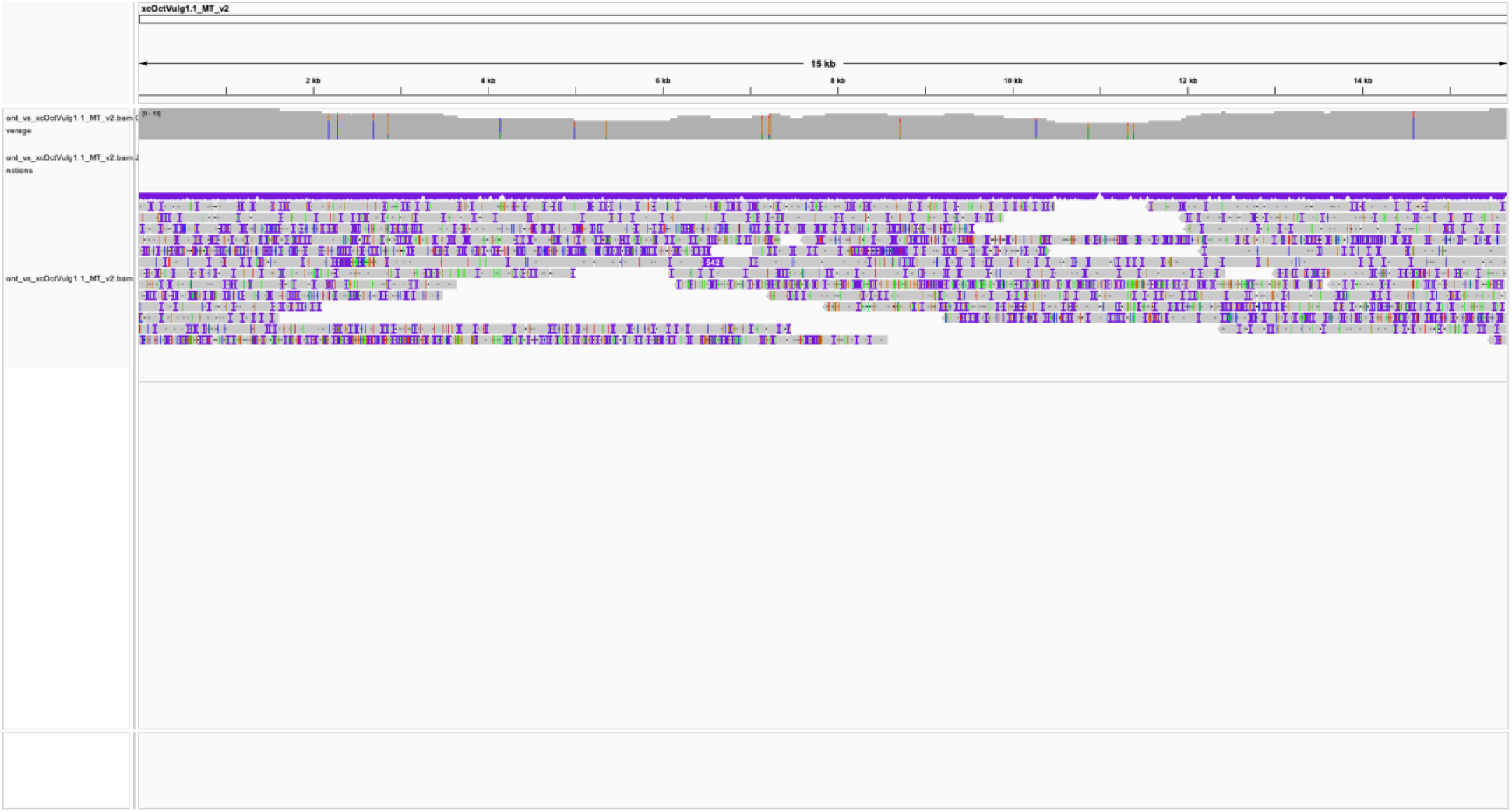
Alignment of ONT reads to the *O. vulgaris* mitochondrial genome. The ONT reads aligned to the mitochondrial genome support the consensus of each position except for 16 nucleotides (see vertical coloured bars along the coverage track).

**Supplementary Figure 6.**
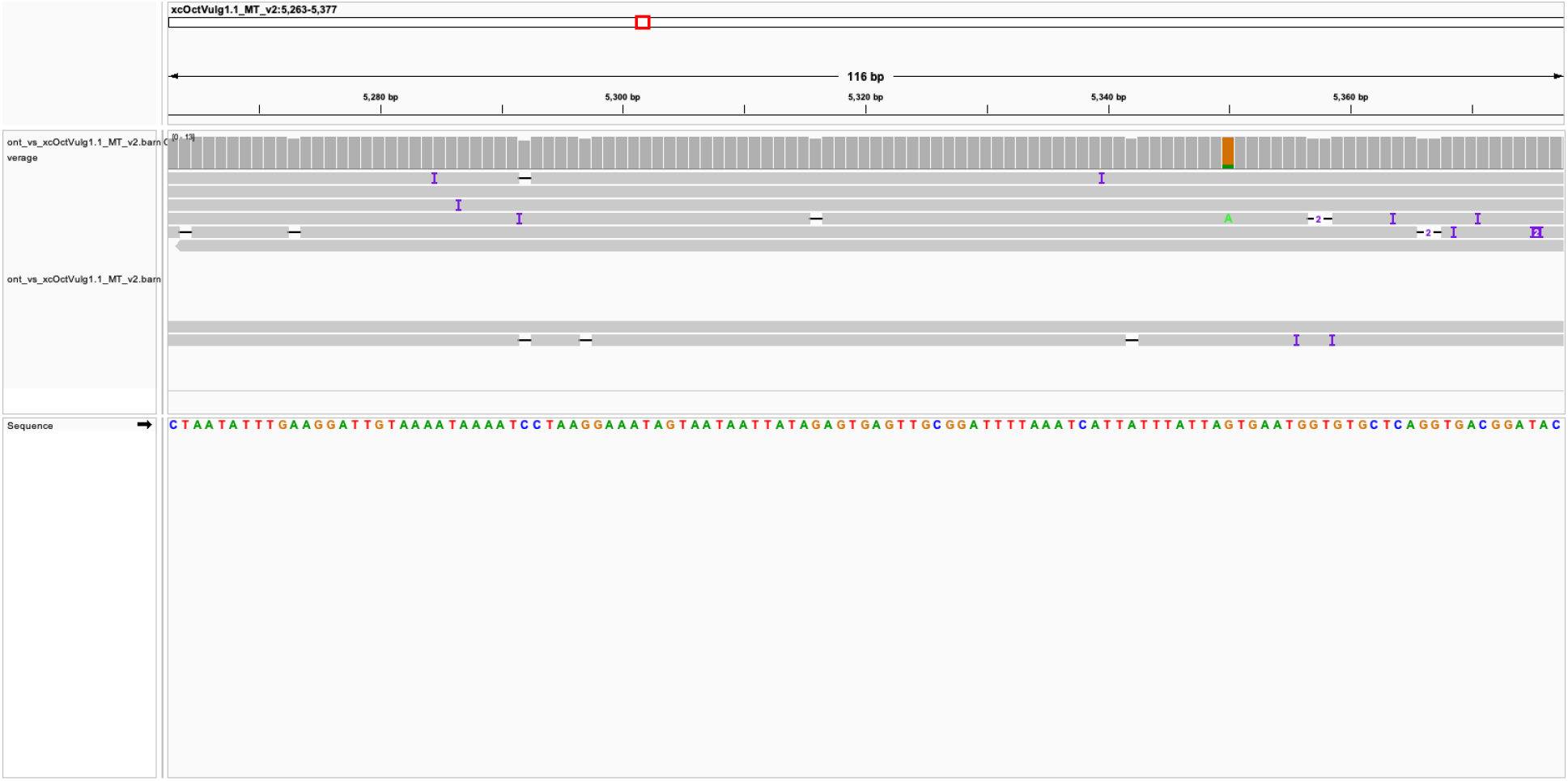
Coverage of ambiguous positions in the mitochondrial genome. The 16 positions that were polymorphic were single-nucleotide differences. The consensus sequence is that with the highest frequency in the reads. One example is shown here, with the G in the mitogenome appearing in most of the reads except one with an A.

## Literature Cited

Adachi K, Ohnishi K, Kuramochi T, Yoshinaga T, Okumura S-I. 2014. Molecular cytogenetic study in *Octopus* (*Amphioctopus*) *areolatus* from Japan. Fish Sci. 80(3):445–450. doi:10.1007/s12562-014-0703-4.

Aho AV, Kernighan BW, Weinberger PJ. 1988. The AWK programming language. Reading, Mass: Addison-Wesley Pub. Co.

[GCF_001194135.2] Albertin CB, Medina-Ruiz S, Mitros T, Schmidbaur H, Sanchez G, Wang ZY, Grimwood J, Rosenthal JJC, Ragsdale CW, Simakov O, et al. 2022. Genome and transcriptome mechanisms driving cephalopod evolution. Nat Commun. 13(1):2427. doi:10.1038/s41467-022-29748-w.

Albertin CB, Simakov O, Mitros T, Wang ZY, Pungor JR, Edsinger-Gonzales E, Brenner S, Ragsdale CW, Rokhsar DS. 2015. The octopus genome and the evolution of cephalopod neural and morphological novelties. Nature. 524(7564):220–224. doi:10.1038/nature14668.

Altschul SF, Gish W, Miller W, Myers EW, Lipman DJ. 1990. Basic local alignment search tool. J Mol Biol. 215(3):403–410. doi:10.1016/S0022-2836(05)80360-2.

Amor MD, Doyle SR, Norman MD, Roura A, Hall NE, Robinson AJ, Leite TS, Strugnell JM. 2019. Genome-wide sequencing uncovers cryptic diversity and mito-nuclear discordance in the *Octopus vulgaris* species complex. bioRxiv 573493. [accessed 2023 Feb 5]. doi:10.1101/573493.

Amor MD, Norman MD, Roura A, Leite TS, Gleadall IG, Reid A, Perales-Raya C, Lu C, Silvey CJ, Vidal EAG, et al. 2017. Morphological assessment of the *Octopus vulgaris* species complex evaluated in light of molecular-based phylogenetic inferences. Zool Scr. 46(3):275–288. doi:10.1111/zsc.12207.

Amor, MD. 2023. Untangling the Octopus vulgaris species complex using a combined genomic and morphological approach [dissertation]. Melbourne: La Trobe University. doi:10.26181/21854307.V1.

Andrews PLR, Darmaillacq A-S, Dennison N, Gleadall IG, Hawkins P, Messenger JB, Osorio D, Smith VJ, Smith JA. 2013. The identification and management of pain, suffering and distress in cephalopods, including anaesthesia, analgesia and humane killing. J Exp Mar Biol Ecol. 447:46–64. doi:10.1016/j.jembe.2013.02.010.

Arbogast BS, Slowinski JB. 1998. Pleistocene speciation and the mitochondrial DNA clock. Science. 282(5396):1955–1955. doi:10.1126/science.282.5396.1955a

Bernt M, Donath A, Jühling F, Externbrink F, Florentz C, Fritzsch G, Pütz J, Middendorf M, Stadler PF. 2013. MITOS: Improved de novo metazoan mitochondrial genome annotation. Mol Phylogenet Evol. 69(2):313–319. doi:10.1016/j.ympev.2012.08.023.

Borrelli L, Gherardi F, Fiorito G. 2006. A Catalogue of Body Patterning in Cephalopoda. 1st ed. Firenze: Firenze University Press (Cataloghi e collezioni). doi:10.36253/88-8453-376-7.

Bredeson JV, Lyons JB, Oniyinde IO, Okereke NR, Kolade O, Nnabue I, Nwadili CO, Hřibová E, Parker M, Nwogha J, et al. 2022. Chromosome evolution and the genetic basis of agronomically important traits in greater yam. Nat Commun. 13(1):2001. doi:10.1038/s41467-022-29114-w.

Buchfink B, Reuter K, Drost H-G. 2021. Sensitive protein alignments at tree-of-life scale using DIAMOND. Nat Methods. 18(4):366–368. doi:10.1038/s41592-021-01101-x.

Cabanettes F, Klopp C. 2018. D-GENIES: dot plot large genomes in an interactive, efficient and simple way. PeerJ. 6:e4958. doi:10.7717/peerj.4958.

Challis R, Richards E, Rajan J, Cochrane G, Blaxter M. 2020. BlobToolKit – Interactive Quality Assessment of Genome Assemblies. G3 GenesGenomesGenetics. 10(4):1361–1374. doi:10.1534/g3.119.400908.

Chapman JA, Ho I, Sunkara S, Luo S, Schroth GP, Rokhsar DS. 2011. Meraculous: De Novo Genome Assembly with Short Paired-End Reads. Salzberg SL, editor. PLoS ONE. 6(8):e23501. doi:10.1371/journal.pone.0023501.

Chiao C-C, Hanlon RT. 2019. Rapid Adaptive Camouflage in Cephalopods. In: Oxford Research Encyclopedia of Neuroscience. Oxford University Press. doi:10.1017/CBO9780511852053.009.

Conesa A, Gotz S, Garcia-Gomez JM, Terol J, Talon M, Robles M. 2005. Blast2GO: a universal tool for annotation, visualization and analysis in functional genomics research. Bioinformatics. 21(18):3674–3676. doi:10.1093/bioinformatics/bti610.

Coombe L, Zhang J, Vandervalk BP, Chu J, Jackman SD, Birol I, Warren RL. 2018. ARKS: chromosome-scale scaffolding of human genome drafts with linked read kmers. BMC Bioinformatics. 19(1):234. doi:10.1186/s12859-018-2243-x.

Dehal P, Boore JL. 2005. Two rounds of whole genome duplication in the ancestral vertebrate. PLoS Biol. 3(10):e314. doi:10.1371/journal.pbio.0030314

Deryckere A, Styfhals R, Elagoz AM, Maes GE, Seuntjens E. 2021. Identification of neural progenitor cells and their progeny reveals long distance migration in the developing octopus brain. eLife. 10:e69161. doi:10.7554/eLife.69161.

Deryckere A, Styfhals R, Vidal EAG, Almansa E, Seuntjens E. 2020. A practical staging atlas to study embryonic development of *Octopus vulgaris* under controlled laboratory conditions. BMC Dev Biol. 20(1):7. doi:10.1186/s12861-020-00212-6.

Dobin A, Davis CA, Schlesinger F, Drenkow J, Zaleski C, Jha S, Batut P, Chaisson M, Gingeras TR. 2013. STAR: ultrafast universal RNA-seq aligner. Bioinformatics. 29(1):15–21. doi:10.1093/bioinformatics/bts635.

Dudchenko O, Batra SS, Omer AD, Nyquist SK, Hoeger M, Durand NC, Shamim MS, Machol I, Lander ES, Aiden AP, et al. 2017. De novo assembly of the *Aedes aegypti* genome using Hi-C yields chromosome-length scaffolds. Science. 356(6333):92–95. doi:10.1126/science.aal3327.

Dudchenko O, Shamim MS, Batra SS, Durand NC, Musial NT, Mostofa R, Pham M, Glenn St Hilaire B, Yao W, Stamenova E, et al. 2018. The Juicebox Assembly Tools module facilitates *de novo* assembly of mammalian genomes with chromosome-length scaffolds for under $1000. bioRxiv 254797. [accessed 2023 Apr 4]. doi:10.1101/254797.

Durand NC, Robinson JT, Shamim MS, Machol I, Mesirov JP, Lander ES, Aiden EL. 2016. Juicebox Provides a Visualization System for Hi-C Contact Maps with Unlimited Zoom. Cell Syst. 3(1):99–101. doi:10.1016/j.cels.2015.07.012.

Fiorito G, Affuso A, Anderson DB, Basil J, Bonnaud L, Botta G, Cole A, D’Angelo L, De Girolamo P, Dennison N, et al. 2014. Cephalopods in neuroscience: regulations, research and the 3Rs. Invert Neurosci. 14(1):13–36. doi:10.1007/s10158-013-0165-x.

Fiorito G, Affuso A, Basil J, Cole A, de Girolamo P, D’Angelo L, Dickel L, Gestal C, Grasso F, Kuba M, et al. 2015. Guidelines for the Care and Welfare of Cephalopods in Research –A consensus based on an initiative by CephRes, FELASA and the Boyd Group. Lab Anim. 49(2_suppl):1–90. doi:10.1177/0023677215580006.

Formenti G, Rhie A, Balacco J, Haase B, Mountcastle J, Fedrigo O, Brown S, Capodiferro MR, Al-Ajli FO, Ambrosini R, et al. 2021. Complete vertebrate mitogenomes reveal widespread repeats and gene duplications. Genome Biol. 22(1):120. doi:10.1186/s13059-021-02336-9.

Gleadall IG. 2016. *Octopus sinensis* d’Orbigny, 1841 (Cephalopoda: Octopodidae): Valid Species Name for the Commercially Valuable East Asian Common Octopus. Species Divers. 21(1):31–42. doi:10.12782/sd.21.1.031.

Gao YM, Natsukari Y. 1990. Karyological Studies on Seven Cephalopods. Venus Jpn J Malacol. 49(2): 126–145. doi:10.18941/venusjjm.49.2_126.

García-Fernández P, Prado-Alvarez M, Nande M, Garcia de la serrana D, Perales-Raya C, Almansa E, Varó I, Gestal C. 2019. Global impact of diet and temperature over aquaculture of *Octopus vulgaris* paralarvae from a transcriptomic approach. Sci Rep. 9(1):10312. doi:10.1038/s41598-019-46492-2.

Guan D, McCarthy SA, Wood J, Howe K, Wang Y, Durbin R. 2020. Identifying and removing haplotypic duplication in primary genome assemblies. Valencia A, editor. Bioinformatics. 36(9):2896–2898. doi:10.1093/bioinformatics/btaa025.

Haas BJ, Salzberg SL, Zhu W, Pertea M, Allen JE, Orvis J, White O, Buell CR, Wortman JR. 2008. Automated eukaryotic gene structure annotation using EVidenceModeler and the Program to Assemble Spliced Alignments. Genome Biol. 9(1):R7. doi:10.1186/gb-2008-9-1-r7.

Hochner B. 2012. An embodied view of octopus neurobiology. Curr Biol. 22(20):R887–R892. doi:10.1016/j.cub.2012.09.001

Hochner B, Shomrat T, Fiorito G. 2006. The octopus: a model for a comparative analysis of the evolution of learning and memory mechanisms. Biol Bull. 210(3):308–317. doi:10.2307/4134567

Hu J, Wang Z, Sun Z, Hu B, Ayoola AO, Liang F, Li J, Sandoval JR, Cooper DN, Ye K, et al. 2023. An efficient error correction and accurate assembly tool for noisy long reads. bioRxiv 2023.03.09.531669. [accessed 2023 Mar 24]. doi:10.1101/2023.03.09.531669.

Inaba A. 1959. Notes on the Chromosomes of Two Species of Octopods (Cephalopoda, Mollusca). Jpn J Genet. 34(5):137–139. doi:10.1266/jjg.34.137.

Iwata H, Gotoh O. 2012. Benchmarking spliced alignment programs including Spaln2, an extended version of Spaln that incorporates additional species-specific features. Nucleic Acids Res. 40(20):e161–e161. doi:10.1093/nar/gks708.

Jackman SD, Coombe L, Chu J, Warren RL, Vandervalk BP, Yeo S, Xue Z, Mohamadi H, Bohlmann J, Jones SJM, et al. 2018. Tigmint: correcting assembly errors using linked reads from large molecules. BMC Bioinformatics. 19(1):393. doi:10.1186/s12859-018-2425-6.

Jiang D, Liu Q, Sun J, Liu S, Fan G, Wang L, Zhang Y, Seim I, An S, Liu X, et al. 2022. The gold-ringed octopus (Amphioctopus fangsiao) genome and cerebral single-nucleus transcriptomes provide insights into the evolution of karyotype and neural novelties. BMC Biol. 20(1):289. doi:10.1186/s12915-022-01500-2.

Jones P, Binns D, Chang H-Y, Fraser M, Li W, McAnulla C, McWilliam H, Maslen J, Mitchell A, Nuka G, et al. 2014. InterProScan 5: genome-scale protein function classification. Bioinformatics. 30(9):1236–1240. doi:10.1093/bioinformatics/btu031.

Kim B-M, Kang S, Ahn D-H, Jung S-H, Rhee H, Yoo JS, Lee J-E, Lee S, Han Y-H, Ryu K-B, et al. 2018 Sep 25. The genome of common long-arm octopus *Octopus minor*. GigaScience. doi:10.1093/gigascience/giy119.

Kolmogorov M, Yuan J, Lin Y, Pevzner PA. 2019. Assembly of long, error-prone reads using repeat graphs. Nat Biotechnol. 37(5):540–546. doi:10.1038/s41587-019-0072-8.

Köster J, Rahmann S. 2012. Snakemake—a scalable bioinformatics workflow engine. Bioinformatics. 28(19):2520–2522. doi:10.1093/bioinformatics/bts480.

Kundu R, Casey J, Sung W-K. 2019. HyPo: Super Fast & Accurate Polisher for Long Read Genome Assemblies. bioRxiv 2019.12.19.882506. doi:10.1101/2019.12.19.882506.

Kurtz S, Phillippy A, Delcher AL, Smoot M, Shumway M, Antonescu C, Salzberg SL. 2004. Versatile and open software for comparing large genomes. Genome Biol. 5(2):R12. doi:10.1186/gb-2004-5-2-r12.

[GCF_006345805.1] Li F, Bian L, Ge J, Han F, Liu Z, Li X, Liu Y, Lin Z, Shi H, Liu C, et al. 2020. Chromosome-level genome assembly of the East Asian common octopus (Octopus sinensis) using PacBio sequencing and Hi-C technology. Mol Ecol Resour. 20(6):1572–1582. doi:10.1111/1755-0998.13216.

Li H. 2013. Aligning sequence reads, clone sequences and assembly contigs with BWA-MEM. doi:10.48550/ARXIV.1303.3997.

Li H. 2018. Minimap2: pairwise alignment for nucleotide sequences. Birol I, editor. Bioinformatics. 34(18):3094–3100. doi:10.1093/bioinformatics/bty191.

Li H, Handsaker B, Wysoker A, Fennell T, Ruan J, Homer N, Marth G, Abecasis G, Durbin R, 1000 Genome Project Data Processing Subgroup. 2009. The Sequence Alignment/Map format and SAMtools. Bioinformatics. 25(16):2078–2079. doi:10.1093/bioinformatics/btp352.

Lomsadze A, Burns PD, Borodovsky M. 2014. Integration of mapped RNA-Seq reads into automatic training of eukaryotic gene finding algorithm. Nucleic Acids Res. 42(15):e119–e119. doi:10.1093/nar/gku557.

Manni M, Berkeley MR, Seppey M, Simão FA, Zdobnov EM. 2021. BUSCO Update: Novel and Streamlined Workflows along with Broader and Deeper Phylogenetic Coverage for Scoring of Eukaryotic, Prokaryotic, and Viral Genomes. Kelley J, editor. Mol Biol Evol. 38(10):4647–4654. doi:10.1093/molbev/msab199.

Mapleson D, Venturini L, Kaithakottil G, Swarbreck D. 2018. Efficient and accurate detection of splice junctions from RNA-seq with Portcullis. GigaScience. 7(12). doi:10.1093/gigascience/giy131.

Marini G, De Sio F, Ponte G, Fiorito G. 2017. Behavioral Analysis of Learning and Memory in Cephalopods ⋆. In: Learning and Memory: A Comprehensive Reference. Elsevier. p. 441–462. doi:10.1016/B978-0-12-809324-5.21024-9.

Marino A, Kizenko A, Wong WY, Ghiselli F, Simakov O. 2022. Repeat Age Decomposition Informs an Ancient Set of Repeats Associated With Coleoid Cephalopod Divergence. Front Genet. 13:793734. doi:10.3389/fgene.2022.793734.

Meyer A, Schartl M. 1999. Gene and genome duplications in vertebrates: the one-to-four (-to-eight in fish) rule and the evolution of novel gene functions. Curr Opin Cell Biol. 11(6):699–704. doi:10.1016/s0955-0674(99)00039-3

Niknafs YS, Pandian B, Iyer HK, Chinnaiyan AM, Iyer MK. 2017. TACO produces robust multisample transcriptome assemblies from RNA-seq. Nat Methods. 14(1):68–70. doi:10.1038/nmeth.4078.

Open2C, Abdennur N, Fudenberg G, Flyamer IM, Galitsyna AA, Goloborodko A, Imakaev M, Venev SV. 2023. Pairtools: from sequencing data to chromosome contacts. bioRxiv 2023.02.13.528389. [accessed 2023 Apr 4]. doi:10.1101/2023.02.13.528389

Pertea M, Pertea GM, Antonescu CM, Chang T-C, Mendell JT, Salzberg SL. 2015. StringTie enables improved reconstruction of a transcriptome from RNA-seq reads. Nat Biotechnol. 33(3):290–295. doi:10.1038/nbt.3122.

Petrosino G, Ponte G, Volpe M, Zarrella I, Ansaloni F, Langella C, Di Cristina G, Finaurini S, Russo MT, Basu S, et al. 2022. Identification of LINE retrotransposons and long non-coding RNAs expressed in the octopus brain. BMC Biol. 20(1):116. doi:10.1186/s12915-022-01303-5.

Ponte G, Chiandetti C, Edelman DB, Imperadore P, Pieroni EM, Fiorito G. 2022. Cephalopod Behavior: From Neural Plasticity to Consciousness. Front Syst Neurosci. 15:787139. doi:10.3389/fnsys.2021.787139.

Ponte G, Taite M, Borrelli L, Tarallo A, Allcock AL, Fiorito G. 2021. Cerebrotypes in Cephalopods: Brain Diversity and Its Correlation With Species Habits, Life History, and Physiological Adaptations. Front Neuroanat. 14:565109. doi:10.3389/fnana.2020.565109.

Rhie A, Walenz BP, Koren S, Phillippy AM. 2020. Merqury: reference-free quality, completeness, and phasing assessment for genome assemblies. Genome Biol. 21(1):245. doi:10.1186/s13059-020-02134-9.

Robinson JT, Thorvaldsdottir H, Turner D, Mesirov JP. 2023. igv.js: an embeddable JavaScript implementation of the Integrative Genomics Viewer (IGV). Alkan C, editor. Bioinformatics. 39(1):btac830. doi:10.1093/bioinformatics/btac830.

Schmidbaur H, Kawaguchi A, Clarence T, Fu X, Hoang OP, Zimmermann B, Ritschard EA, Weissenbacher A, Foster JS, Nyholm SV, et al. 2022. Emergence of novel cephalopod gene regulation and expression through large-scale genome reorganization. Nat Commun. 13(1):2172. doi:10.1038/s41467-022-29694-7.

Shigeno S, Andrews PLR, Ponte G, Fiorito G. 2018. Cephalopod Brains: An Overview of Current Knowledge to Facilitate Comparison With Vertebrates. Front Physiol. 9:952. doi:10.3389/fphys.2018.00952.

Smit AFA, Hubley R, Green P. 2013-2015. RepeatMasker Open-4.0. http://www.repeatmasker.org.

Stanke M, Schöffmann O, Morgenstern B, Waack S. 2006. Gene prediction in eukaryotes with a generalized hidden Markov model that uses hints from external sources. BMC Bioinformatics. 7(1):62. doi:10.1186/1471-2105-7-62.

Styfhals R, Zolotarov G, Hulselmans G, Spanier KI, Poovathingal S, Elagoz AM, De Winter S, Deryckere A, Rajewsky N, Ponte G, et al. 2022. Cell type diversity in a developing octopus brain. Nat Commun. 13(1):7392. doi:10.1038/s41467-022-35198-1.

Taite M, Fernández-Álvarez FÁ, Braid HE, Bush SL, Bolstad K, Drewery J, Mills S, Strugnell JM, Vecchione M, Villanueva R, et al. 2023. Genome skimming elucidates the evolutionary history of Octopoda. Mol Phylogenet Evol. 182:107729. doi:10.1016/j.ympev.2023.107729.

Vara C, Paytuví-Gallart A, Cuartero Y, Álvarez-González L, Marín-Gual L, Garcia F, Florit-Sabater B, Capilla L, Sanchéz-Guillén RA, Sarrate Z, et al. 2021. The impact of chromosomal fusions on 3D genome folding and recombination in the germ line. Nat Commun. 12(1):2981. doi:10.1038/s41467-021-23270-1.

Vitturi R, Rasotto MB, Farinella-Ferruzza N. 1982. The chromosomes of 16 molluscan species. Bolletino Zool. 49(1–2):61–71. doi:10.1080/11250008209439373.

Wang J, Zheng X. 2017. Comparison of the genetic relationship between nine Cephalopod species based on cluster analysis of karyotype evolutionary distance. Comp Cytogenet. 11(3):477–494. doi:10.3897/compcytogen.v11i3.12752.

Wang ZY, Ragsdale CW. 2019. Cephalopod Nervous System Organization. In: Oxford Research Encyclopedia of Neuroscience. Oxford University Press. doi:10.1093/acrefore/9780190264086.013.181

Warren RL, Yang C, Vandervalk BP, Behsaz B, Lagman A, Jones SJM, Birol I. 2015. LINKS: Scalable, alignment-free scaffolding of draft genomes with long reads. GigaScience. 4(1):35. doi:10.1186/s13742-015-0076-3.

Weisenfeld NI, Kumar V, Shah P, Church DM, Jaffe DB. 2017. Direct determination of diploid genome sequences. Genome Res. 27(5):757–767. doi:10.1101/gr.214874.116.

Whitelaw BL, Jones DB, Guppy J, Morse P, Strugnell JM, Cooke IR, Zenger K. 2022. High-Density Genetic Linkage Map of the Southern Blue-ringed Octopus (Octopodidae: Hapalochlaena maculosa). Diversity. 14(12):1068. doi:10.3390/d14121068.

Yokobori S-i. 2004. Long-Term Conservation of Six Duplicated Structural Genes in Cephalopod Mitochondrial Genomes. Mol Biol Evol. 21(11):2034–2046. doi:10.1093/molbev/msh227.

Young JZ. 1964. A Model of the Brain. Oxford University Press, USA.

Zarrella I, Herten K, Maes GE, Tai S, Yang M, Seuntjens E, Ritschard EA, Zach M, Styfhals R, Sanges R, et al. 2019. The survey and reference assisted assembly of the *Octopus vulgaris* genome. Sci Data. 6(1):13. doi:10.1038/s41597-019-0017-6.

Zhang H, Song L, Wang X, Cheng H, Wang C, Meyer CA, Liu T, Tang M, Aluru S, Yue F, et al. 2021. Fast alignment and preprocessing of chromatin profiles with Chromap. Nat Commun. 12(1):6566. doi:10.1038/s41467-021-26865-w.

Zhou C, McCarthy SA, Durbin R. 2023. YaHS: yet another Hi-C scaffolding tool. Alkan C, editor. Bioinformatics. 39(1):btac808. doi:10.1093/bioinformatics/btac808.

Zolotarov G, Fromm B, Legnini I, Ayoub S, Polese G, Maselli V, Chabot PJ, Vinther J, Styfhals R, Seuntjens E, et al. 2022. MicroRNAs are deeply linked to the emergence of the complex octopus brain. Sci Adv. 8(47):eadd9938. doi:10.1126/sciadv.add9938.

